# Study of the DiI method accuracy on the example of the afferent connections of the Habenula in P6 F1-C57Bl6 / CBA mice

**DOI:** 10.1101/277541

**Authors:** A Klepukov, V.P Apryshko, S.A. Yakovenko

## Abstract

DiI (1,1’-dioctadecyl-3,3,3’3’-tetramethylindocarbocyanine perchlorate) is a lipophilic carbocyanine stain diffusing in the plasma membrane and coloring the cell as a whole.

The DiI injection method is an alternative to stereotaxis in the study of the neural connections of higher vertebrates. However, despite a number of studies performed to identify the neural connections of embryos and fetuses of different animals, the question of the accuracy of the DiI method remains open. The main goal of this study is to investigate the accuracy of the DiI method by examining the afferent Habenula (Hb) connections in P6 F1-C57Bl6/CBA mice. Hb is a paired structure of the intermediate brain, consisting of two nuclei - the medial (MHb) and lateral (LHb). At P6, the main Hb tracts are already formed but not yet myelinated. To estimate the accuracy of the method, two groups of cases with different types of application on different Hb nuclei were compared: bilateral applications of several DiI crystals - group 1 (26 applications); Unilateral applications of a single crystal DiI - control group 2 (7 applications). It is known that MHb is strictly innervated by Triangular septal nuclei (TS) and Septofimbrial septal nuclei (SFi); LHb is strictly innervated by the Lateral preopteic area (LPA). Any deviation from this scheme is a false positive result. When applying the marker to different Hb nuclei in group 1, the number of successful cases without deviations was -0%; in group 2 (control) - 28.57%. Unilateral applications of a single DiI crystal are more accurate. However, there are still a number of unavoidable problems, such as the fragmentation or loss of a single DiI crystal during application and the super-strong diffusion of the marker at the application site. Given this, the overall accuracy of the DiI injection method, even in the form of a single crystal, remains low.

## 1 INTRODUCTION

Generally, microscopy research methods of intracerebral connections in higher vertebrates are universal. The overwhelming majority of these methods are based on the sterotactic injections of various markers in the desired brain structure. However, stereotaxis has a number of significant limitations, for example, it is applicable only to live brain tissue, it is not applicable to the study of human nerve pathways, the high cost of stereotaxic apparatus and the limitations imposed by the postoperative period make this method of research extremely expensive (Blackstad TW, Heimer L, Mugnaini E. 1981). All these limitations logically lead to the search for alternative research methods of nerve connections, avoiding stereotaxis. And such a method has been found - this is a method of intracerebral connections studying by injecting lipophilic carbocyanine dyes - such as DiI. (Honig M, Hume R. 1986). DiI (1,1’-dioctadecyl- 3,3,3’3’-tetramethylindocarbocyanine perchlorate) passively diffuses in the lipid bilayer of the plasma membrane, staining the whole cell with all its endings. The DiI connections research method has a number of undeniable advantages over stereotaxis, but also a number of significant shortcomings (Molnar Z et al., 2006) and their number is comparable (Table 1). This significantly limits the use of the DiI method in the research study of neural connections of higher vertebrates, which has now virtually come to naught, since most of the original work on the study of intracerebral connections by this method was carried out back in the 1990s. However, the potential of this method is evidently not fulfilled enough, especially given that DiI is able to color neurons and neural tracts anywhere in the brain, including that it is of hard access for stereotaxis. Despite the fact that there is a number of works devoted to the study of the of neural connections formation of various brain structures in embryos and fetuses of rats or mice, none of these studies raised the accuracy of the corresponding nerve connections detection with DiI as the main question. A small number of works performed with the use of the DiI method and own preliminary studies (Klepukov A, Makarenko IG. 2013) indicate an imperfection of the methodical part and low accuracy of this neural connections research method.

**Table 1.** Comparison of advantages and disadvantages of DiI as a tracer for the study of intracerebral connections.

| Advantages | Disadvantages |
| --- | --- |
| DiI Detects neural connections in both living and fixed nerve tissue | The long diffusion time of DiI along the neuron membrane. It takes months |
| DiI Detects neural connections of any species of higher vertebrates, does not require detailed stereotaxic atlases | The DiI diffusion distance along the membrane is limited by a distance of 4 mm for any animal species. |
| DiI Detects neural connections of fetuses and neonatal animals, as well as human embryos. | DiI Does not work on myelinated neural tissue. |
| DiI Paint a neuron as a whole | DiI It does not allow to clearly differentiate the axon and dendrites, which greatly complicates the analysis of data |
| DiI Does not require an expensive stereotaxic installation, since the application of the marker is carried out directly into the desired structure on the brain slice | The extensive diffusion of DiI at the site of application often leads to false positive results |

The aim of this paper is to investigate the accuracy of the DiI method and to improve the methodological part, since this may possibly contribute to a new revival of this method. The best way to investigate DiI injection for accuracy, it to test already known brain structures, whose connections are well described, for example, on Habenula (Hb). Hb is a paired structure of the intermediate brain, its afferent connections are described in detail in the literature both for rats (Herkenham M, Nauta WJ, 1977) and partially for mice (Qin C, Luo M. 2009; Broms J et al., 2017). It is known that Medial Habenular nuclei (MHb) is innervated by Septum nuclei, such as Triangular septal nuclei (TS) and Septofimbrial septal nuclei (SFi); Lateral Habenular nuclei is innervated by the Lateral pteoptic area (LPA). The Hb nucleus is an ideal candidate as an indicator of the depositions accuracy, since any deviations in the results from what is described in the literature will be correctly interpreted precisely as the result of inaccurate application of the marker or its excessive diffusion at the application site (false positive result). Given this, by the ratio of the successful cases number (with the similar results described in the literature for the distribution of neurons after the injection of DiI into different Hb nuclei) to failed cases (with a false positive result), it is possible to understand how accurately DiI identifies structures innervating different Hb nuclei.

This study was performed on hybrids of F1-C57Bl6/CBA mice of the sixth day of postnatal development (P6), by this time the main neural tracts have already been formed, but still unmyelinated (Jacobson S. 1963). This is important, since DiI well reveals only the unmyelinated nerve pathways (Molnar Z, Blakemore C. 1995). In this study, only one type of marker application was used: in the form of crystals / crystal. This method of application is not unique, but it is the most common in the study of neural connections by the DiI method (Molnar Z et al., 2006).

## 2 METHODS

1. *Nomenclature*. In this article we will use the nomenclature (Table 2) *of Franklin and Paxinos (2008).*
2. *Animals*. In this study, the Bioethics commission of the Faculty of Biology of Moscow State University (Conclusion 56 of 9.03.2017) has approved all procedures on animals. *Such procedures on animals as perfusion and euthanasia were carried out at the SRI of the Mitengineering of the MSU in accordance with the Directive 2010/63 / EU of the European Parliament and the Council of the European Union for the protection of animals used for scientific purposes.* All the work was carried out on mice - hybrids F1 - C57Bl6/CBA. The total number of animals was 20. The birthday of the mice was considered a zero day of postnatal development (P0), all procedures were performed on mice on the seventh day of postnatal development (designated as P6).
3. *Procedures performed on animals.* All mice (P6) were given intraperitoneal solutions of zoletil 100 (Vibrac) in saline (0.9% NaCl) at the rate of 10 mg per 1 kg of body weight before perfusion. Then mice were subsequently perfused with saline solution (0.9% NaCl) through the left ventricle of the heart to remove blood from the blood vessels. Thereafter, the mice were decapitated and the heads were placed in 4% PAF on 0.1 M PB (pH 7.4) for up to one month term at room temperature for postfixation. After postfixation, the brain was removed from the skull and stored in 4% PAF until the application of DiI crystals.
4. Search for the desired brain level and application of crystals / crystal marker: 1,1’-dioctadecyl- 3,3,3’, 3’-tetramethyl-indocarbocyanine perchlorate (DiI). The crystal / DiI crystal was applied exclusively to the Hb nuclei. Before applying the crystals / crystal, it was necessary to accurately find the level of the brain on the coronary section, corresponding to the level of Hb. To do this, the brain was placed into a warm (45-55C) solution of 5% agar-agar, which after cooling formed a block of the desired shape around the brain. Further, this agar-agar block was placed in a Campden instruments HA 752 bath filled with saline (0.9% NaCl) and sequentially sliced from the caudal side of the brain to a level corresponding to Hb under the control of a stereo microscope (AmScope 07X-45X). After the desired brain level corresponding to Hb was found, the agar-agar block into which the brain was enclosed was extracted outside from the saline solution. The surface of the slice was carefully dried with filter paper. Further, agar agar blocks with the brain were divided into two groups of samples according to the type of application of DiI crystals / crystal. In the first group of samples (group 1; 13 cases), several DiI crystals (the diameter of each crystal is not more than 50 µm) were bilaterally applied to one of the Hb nuclei of each hemisphere of the brain. In the second group of samples (group 2 - control group, 7 cases), a single DiI crystal (crystal diameter not more than 50 µm) was applied strictly unilaterally to one of the Hb nuclei of one hemisphere of the brain. The application of DiI crystals / crystals to Hb was carried out under the control of a stereo microscope (AmScope 07X-45X) equipped with a course micromanipulator (Narishige C-1). A glass needle with an angle of 35 (K-PZD-1035 COOK MEDICAL) was placed on the micromanipulator, then the needle was dropped into the substrate, dotted with DiI crystals, which easily settled on the glass surface of the needle due to adhesive forces. If the DiI crystals formed small clusters, then on the needle they settled in a cluster too, so hereafter this cluster of crystals was used for deposition in Hb (group 1). A single DiI crystal could be captured when it lay apart from all the others, and this crystal was used for deposition in Hb (group 2). The injection of the DiI crystals /crystal itself was carried out by puncturing the surface of the brain slice into one of the Hb nuclei. After the DiI injection, the agar-agar block with the brain was neatly placed in 4% PAF for 0.1 M PB for a period of 1.5 to 3 months. This time was necessary for the diffusion of the marker along the nerve endings. The samples were stored at the room temperature in the dark.
5. *Slicing the material.* The brain in the agar-agar block was placed in a bath filled with saline solution (0.9% NaCl) and subsequently sliced on a vibroslicer (Campden instruments HA752) in a rostro-caudal direction. The thickness of the sections was 150 µm. The slices were placed on L-polilysin-coated glasses (Thermo) with a thin hair brush, lightly dried and placed into a solution of Mowiol 4-88 (Kremer), prepared on glycine-alkaline buffer. The finished specimens were stored at a temperature of + 25 ° C for a week.
6. Specimen analysis. The specimens were examined in a fluorescence microscope (Altami LUM2) equipped with a green filter (Excitation: 460 nm, Emission: 550 nm)) to detect the orange-red fluorescence specific to DiI. The place of application was evaluated in transmitted light. After evaluating the spread of the marker at the application site, a series of brain sections were sequentially looked through, the distribution of stained DiI nerve fibers and neurons in LPA, TS and SFi was described. Photographing of specimens was carried out with a digital camera (Bioimager BIC-C30) and a special computer program for shooting (ToupTek). Computer program Photoshop CS3 (Adobe, USA) was used for processing and editing of digital images and for illustration of the results. An atlas of the developing mouse brain was used to identify the departments and nuclei of the brain, (Paxinos et al., 2007).

**Table 2.** Abbreviations

| Notation | Structure |
| --- | --- |
| BST | Bed nucleus of stria terminalis |
| Hb | Habenula |
| HDB | Diagonal band, horizontal limb |
| EPN | Entopeduncular nucleus |
| LH | Lateral hypothalamic area |
| LHb | Lateral habenular nucleus |
| LHbM | Lateral habenular nucleus, medial part |
| LHbL | Lateral habenular nucleus, lateral part |
| LPA | Lateral preoptic area |
| MHb | Medial habenular nucleus |
| SFi | Septofimbrial nucleus |
| sm | stria medullaris |
| TS | Triangular septal nucleus |

## 3 RESULTS

### 3.1 General comments on the interpretation of the results

Due to the fact that DiI diffuses in the neuron membrane, both in the retrograde and anterograde directions, initially a special planning of the experiment is required for the correct interpretation of the obtained data. It is known (from the literature data) that in mice and rats MHb are strictly innervated by caudal cores of the septum - TS and SFi; LHb is strictly innervated by such nuclei as LPA. Cross-links between adjacent nuclei of MHb and LHb are not described; connections between Hb nuclei of different hemispheres of the brain are not described. Thus, if neurons are detected in LPA when DiI is applied to MHb, then this indicates either the diffusion of the marker between neighboring Hb nuclei (if applied accurately to one core) or the inaccurate application of the marker in Hb (in two nuclei simultaneously). In the event if neurons in TS or SFi are detected when DiI is applied to LHbM or LHbL, the causes are the same as described above. To assess the accuracy of the application of DiI to the presence of labeled bodies of neurons, those structures of the brain that are affiliated with one Hb nucleus, should not be innervated by another nucleus of Hb with a deposited marker. The general scheme of the logic of the experiment is shown in Fig. 1 (A, B, C, D).

**Fig.1.**
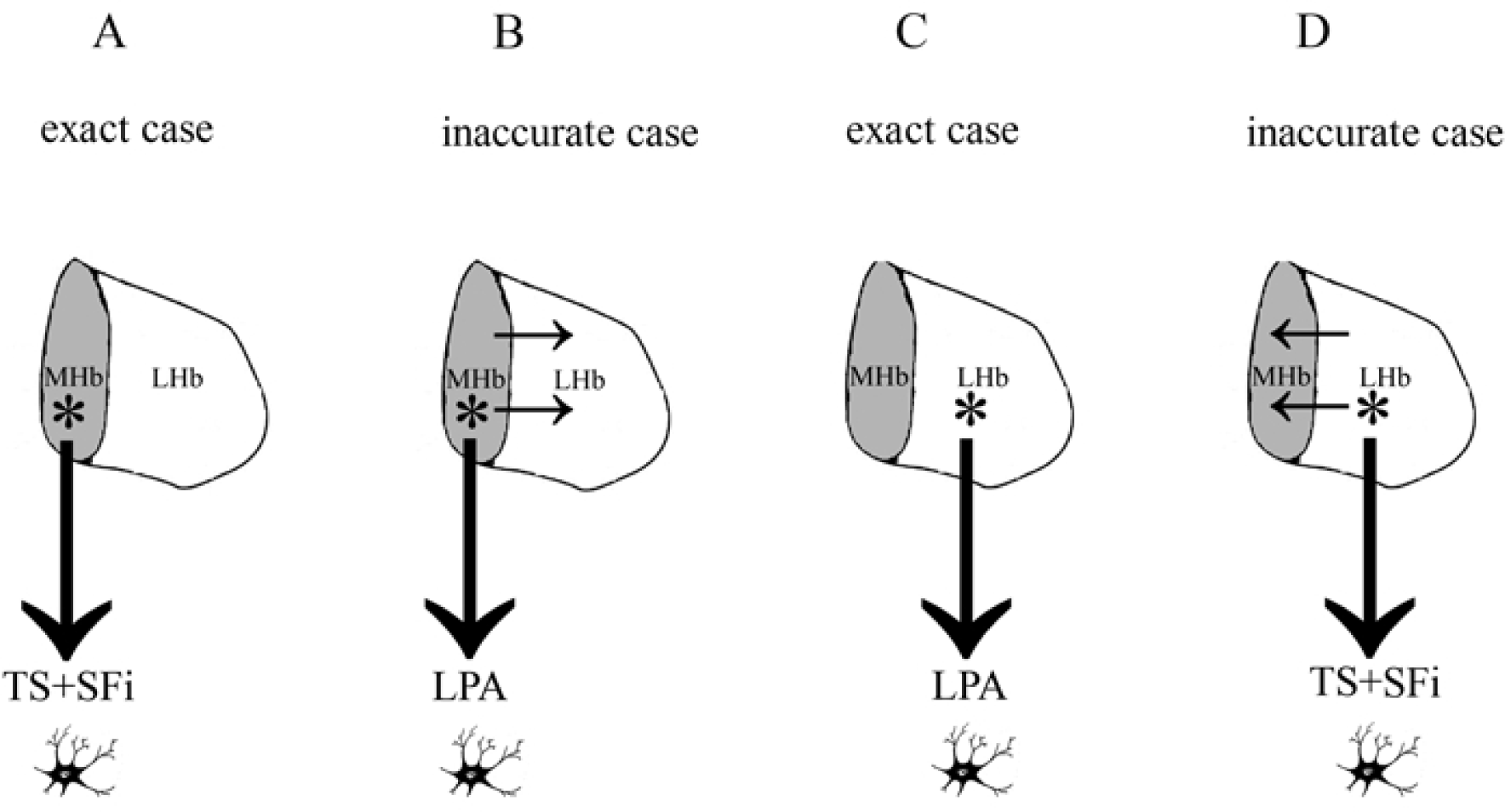
The general scheme ofthe logic of the experiment, which explains how to test Dil for the accuracy ofidentifying intracerebral connections. A - The application of Oil to MHb (marked with an asterisk) strictly reveals the neurons in the caudal nuclei of the septum - TS and SFi, and this is the exact case (corresponding to literature data). B - Application of Oil to MHb (marked with an asterisk) reveals neurons in the LPA - this indicates either the marker diffusion between adjacent Hb nuclei (marked with a horizontal arrow) or the inaccurate marking in Hb (in two nuceli at the same time), it is an inaccurate case. C - The application ofDil to LHb (marked with an asterisk) strictly reveals the neurons in the LPA, and this is an accurate case (corresponding to literature data). D - The application ofDil to LHb (marked with an asterisk) reveals neurons in TS or SF-this indicates either the diffusion of the marker between adjacent Hb nuclei (indicated by the horizontal arrow) or an inaccurate application of the marker in Hb (in two nuceli at the same time) is an inaccurate case.

In this study, the application of DiI to different Hb nuclei was performed in a puncture on the surface of the brain, cut coronary, this provided free access to different sites of Hb - either MHb, or LHbM or LHbL. The boundary between LHbM and LHbL was conventionally carried out along the midline separating the LHb core into two equal parts. Mostly applications were made at the medial level of Hb along the rostro-caudal axis corresponding to a distance of about 600 µm from the start of sm to the site of application (twelve cases, Tables 3,4,5,6).

**Table 3.** Bilateral application of several DiI crystals on MHb - four cases, eight applications. All applications are inaccurate since the marker covered both the MHb and LHb nuceli.

| Case | Place of application<br>(Left hemisphere) | Presence of<br>neurons in<br>LPA | Place of<br>application<br>(Right<br>hemisphere) | Presence of<br>neurons in<br>LPA | Level of<br>application<br>site along the<br>rostro-caudal<br>axis |
| --- | --- | --- | --- | --- | --- |
| P1 | Extensive<br>concentrated<br>application on MHb<br>+ LHb | Reliable<br>Analysis is<br>impossible | Extensive<br>scattered<br>application<br>on<br>MHb + LHb | Reliable<br>Analysis is<br>impossible | Cauda |
| P6 | Extensive<br>concentrated<br>application on MHb<br>+ LHb | Reliable<br>Analysis is<br>impossible | Extensive<br>scattered<br>application<br>on<br>MHb + LHb | Reliable<br>Analysis is<br>impossible | Mid |
| P9 | Extensive<br>concentrated<br>application on MHb<br>+ LHb | Reliable<br>Analysis is<br>impossible | Extensive<br>scattered<br>application<br>on<br>MHb + LHb | Reliable<br>Analysis is<br>impossible | Cauda |
| P13 | Local application<br>On MHb + LHbM | Reliable<br>Analysis is<br>impossible | Local<br>application<br>On MHb +<br>LHbM | Reliable<br>Analysis is<br>impossible | Mid |

**Table 4.**
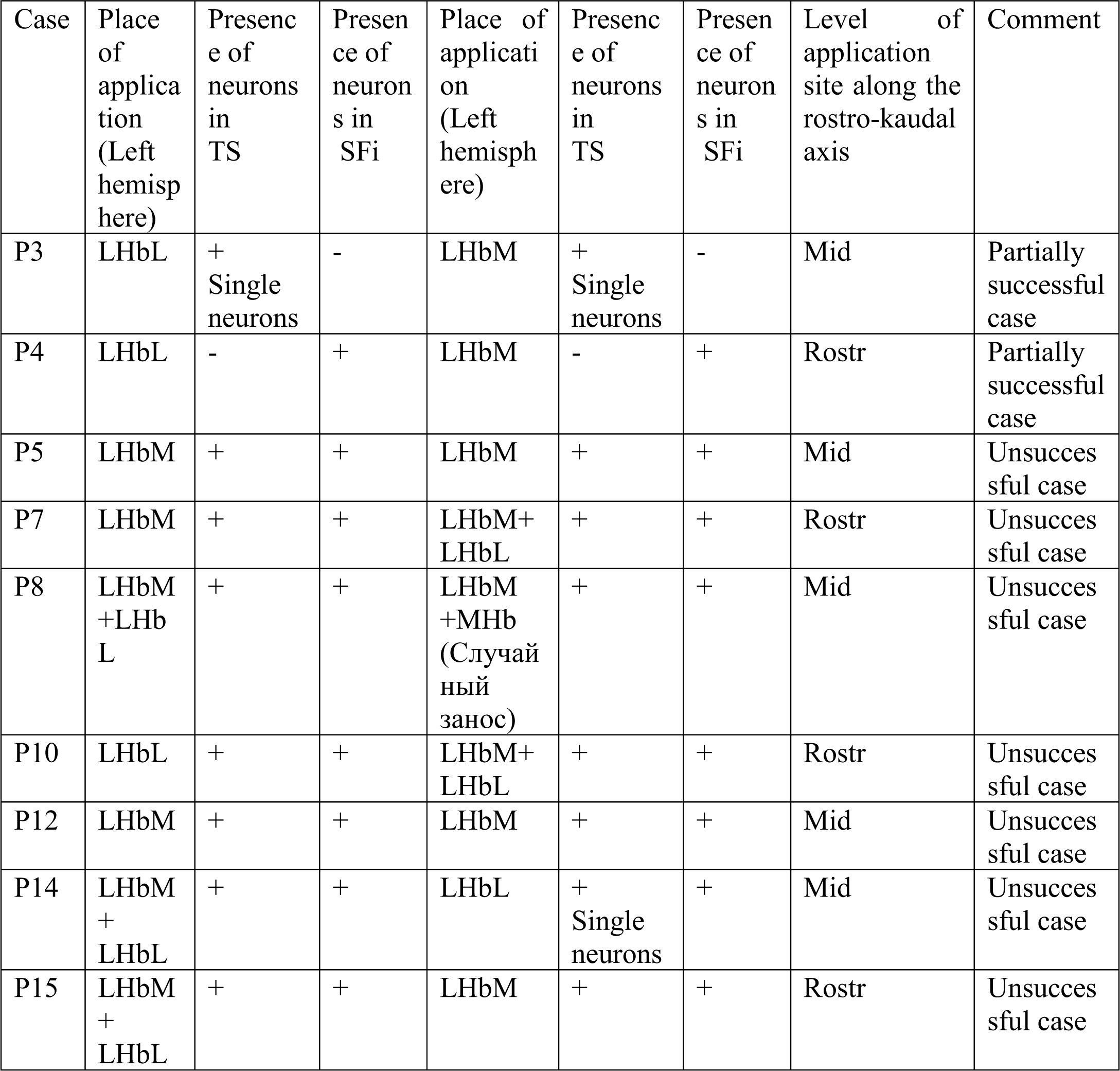
Bilateral application of several DiI crystals on LHb - nine cases. Six DiI applications were made to LHbM, three DiI applications were made to LHbL. The remaining nine applications were inaccurate and affected both parts of the LHb. In one application (P6-8 on the right), a part of the marker got in MHb – a random marker skid.

**Table 5.** Unilateral application of a single DiI crystal to MHb - three cases.

| Case | Place of application | LPA | LHb core of the same hemisphere of the brain | MHb core of the same hemisphere of the brain | Level of application site along the rostro-kaudal axis | Comment |
| --- | --- | --- | --- | --- | --- | --- |
| P6-16 | MHb | +<br>Single neuron | + | + | Mid | The crystal fractured in the tissue when applied |
| P6-11 | MHb | - | + | + | Rostr | Nominally successful result |
| P6-17 | MHb | + | + | + | Mid | The crystal fractured in the tissue when applied |

**Table 6.**
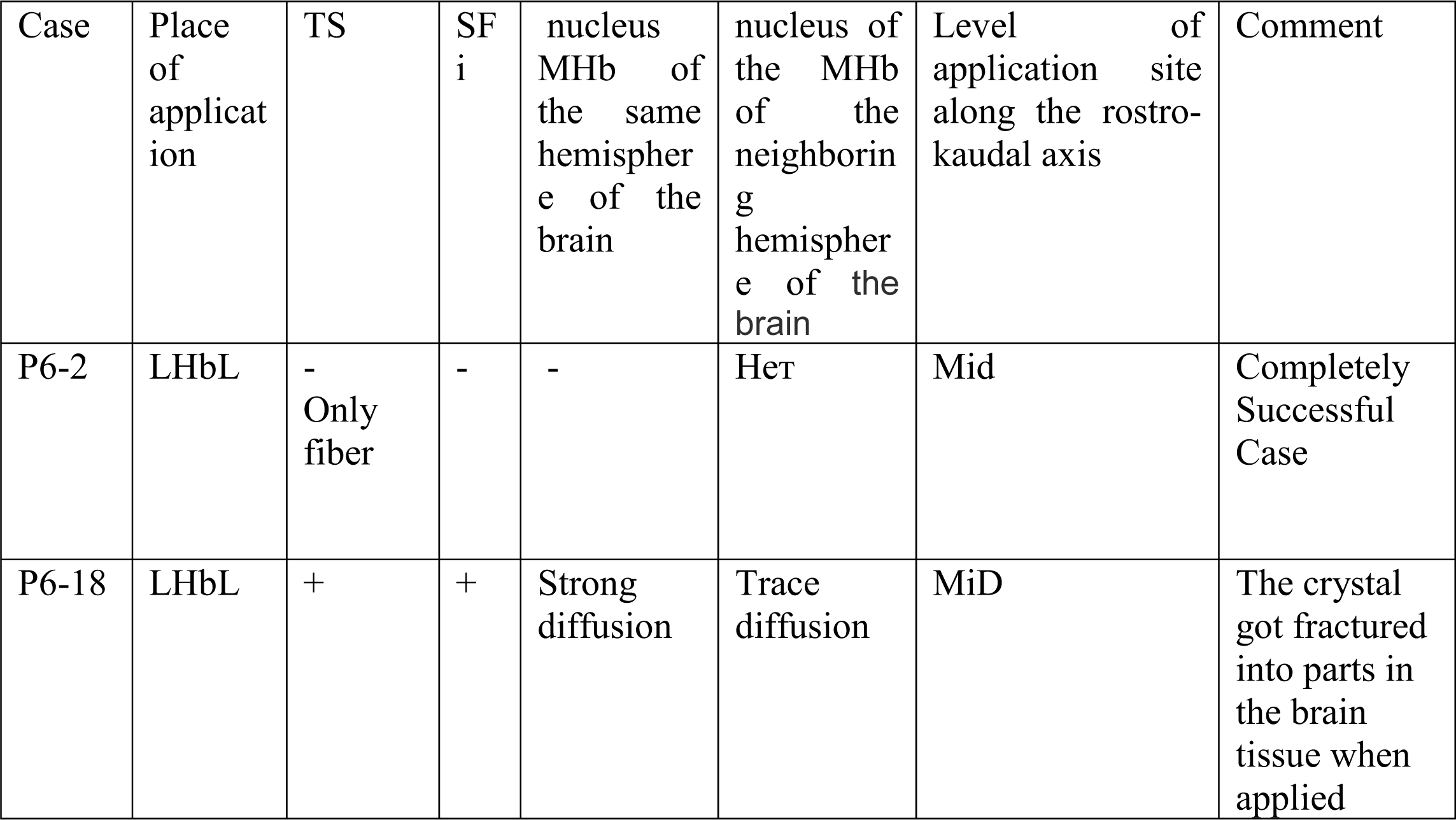
Unilateral application of a single DiI crystal to LHb - two cases.

Some applications were made on the rostral level of Hb, corresponding to a distance of about 300 µm from the start of sm to the site of application (four cases, Tables 3,4,5,6). In two cases, bilateral application of DiI was done at the caudal level of Hb, corresponding to a distance of 900 µm from sm. (Table 3). The total length of Hb was assumed to be about 1200 µm (Paxinos G et al., 2007).

To test the effect of the number of DiI crystals on the accuracy of the application, two groups of brain samples differing in number of deposited crystals (several or one) were used,

### 3.2 Two groups of brain samples with a different strategy of application DiI to Hb

In the first group (group 1), several small crystals of DiI were bilaterally applied into one of the Hb nuclei (MHb, LHbM or LHbL). The total number of cases is thirteen, in total twenty-six DiI applications, twelve successful accurate applications of DiI into one nucleus (Table 4).

The presence of several crystals at the place of application guaranteed that the quantity of the marker are sufficient for the diffusion to be successful. However, there is a risk that the application will be too large and cover both nuceli at the same time MHb and LHb, or both parts of the nucleus in the case of LHb, and this can lead to initially unreliable results. Part of the cases for this reason were completely or partially unsuccessful, depending on whether inaccurate applications were made in one or two hemispheres simultaneously (Table 3, 4).

In the second group (group 2 - control group), a single small DiI crystal was applied to one of the Hb nuclei (MHb or LHb) strictly unilaterally. The total number of cases is seven, and number of successful cases with accurate application is five (Table 5, 6). Unilateral application of a single crystal potentially increased the number of used animals compared to bilateral applications, since DiI applications were not performed in two Hb nuclei of different hemispheres, but into one Hb nucleus in one hemisphere. In addition, the probability of a weak fixation of the DiI crystal increased the risk of lack of results. Two samples of the brain - P6-19 and P6-20 - for this reason were lost and were not analyzed. Besides, there was a risk that a single DiI crystal might simply not be enough for successful diffusion. However, the unilateral application of a single crystal made it possible to evaluate the diffusion of the marker between Hb nuclei from different hemispheres of the brain.

### 3.3 Analysis of places of application

The study of each brain began with sections containing the place of application of the crystals / DiI crystal, followed by the analysis of the distribution of the marker in the area of its application. The area of the marker distribution in fluorescent light was determined by the bright glow of the labeled fibers and neurosomes, and the luminescence intensity was so high at the site of application that it was often difficult to determine the location of the crystals / DiI crystal in the structure under study. Analysis of the application site using an ultraviolet filter is also not suitable, as it has not always been possible to achieve sufficient contrast of the image.

For these reasons, in order to clarify the location of the crystals / DiI crystal and the degree of diffusion of DiI at the sites of application, the evaluation was carried out in a transmitted light. In all cases, undissolved crystals were present in the tissue, which were clearly visible in transmitted light. The diffusion area of DiI at the place of application was determined visually by the presence of a distinctive pinkish coloration in the structure under study, which is well detected in transmitted light (uncoloured tissue is colorless). The difference in the size of the application was not originally planned in the experiment, the size of the application was determined post factum.

### 3.4 Bilateral applications of several DiI crystals - group 1

1. The application of several DiI crystals to MHb - four cases, eight depositions (Table 3, Fig.2.A, B, C, D) in one more case (P6-8, application on the right) when the marker is placed on the LHbM part of the crystals DiI fall on MHb (illustration 3.B). All eight applications of DiI crystals were inaccurate. In the case of P6-1 (application to the left), P6-6 (application to the left), P6-9 (application to the left), the DiI application was too large and extensive and captured both MHb and LHb nuclei. In all three of these cases, applications on the right side were vise-versa small and excessively scattered in area and also affected both MHb and LHb nuclei. In the case of P6-13, the application to MHb were more pointlike, but an insignificant part of the DiI crystals accidentally fell in LHbM in both hemispheres (picture 2D). A reliable analysis of the results of neuronal distribution in the LPA in all these four cases is impossible since none of the applications can be considered accurate.
2. The deposition of several DiI crystals on LHbM - six cases, eight precise depositions in the medial part of the LHb nucleus (Table 4; Fig. 3A, B, E, G; Fig. 4A, B). Despite the fact that the application of DiI crystals was made exactly to the LHbM nucleus, the degree of diffusion of the marker outside the place of application varied greatly. The key factor in these applications was whether the diffusion of the marker spread to the neighboring MHb nucleus. In transmitted light in two applications (P6-3 on the right, P6-4 on the right) out of six this kind of diffusion in MHb was almost not detected, both applications were very local (pic.4.A, B). In other cases, the marker propagated to neighboring nuceli, both in MHb and LHbL.
3. The application of several DiI crystals to LHbL - three cases, four precise applications directly into the lateral part of the LHb nucleus (Table 4; Fig.3.D, F; Fig. 4A, B). As in the previous case, the key factor in these applications was also the absence of diffusion in the neighboring MHb nucleus. In one case out of four (P6-3 on the left), undesirable diffusion in MHb was practically not fixed, because the application was very local. The remaining two applications, on the contrary, were distinguished by a larger area and more pronounced strong diffusion of DiI beyond the nucleus boundaries.
4. The application of several DiI crystals on LHbM and LHbM - five cases, six depositions (Table 4; Fig. B, C, D, F, G). All these cases are considered inaccurate. Those disposition was considered inaccurate, in which undissolved Dill crystal is different parts of one nucleus– LHbM И LHbL – were evidently fixed in the transmitted light. In one case (P6-8, application to the left), the presence of small DiI crystals in MHb (random drift) was recorded.

**Fig.2.**
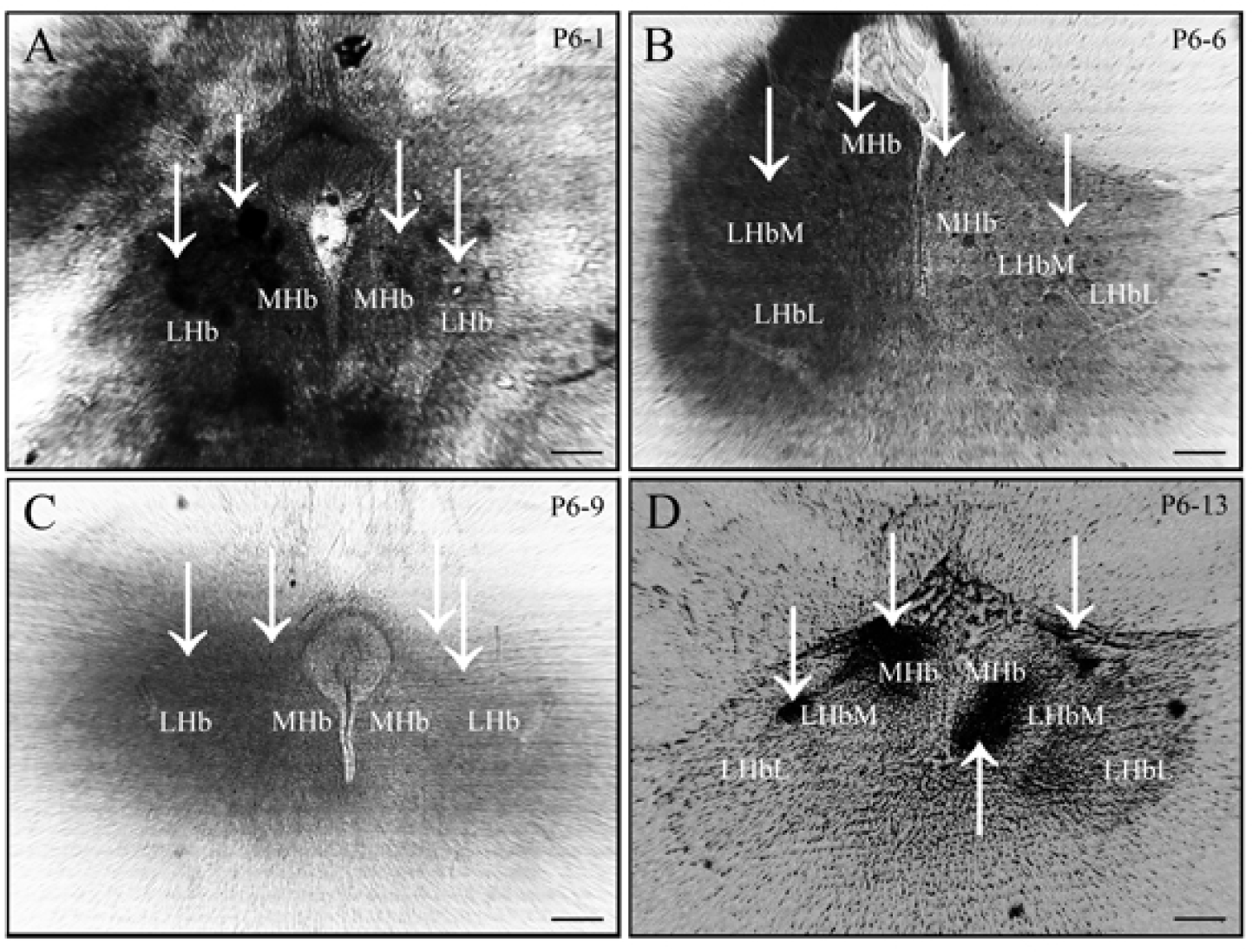
A, B, C, D - Bilateral sites for applying several Oil crystals to MHb, four cases (P6-1, P6-6, P6-9, P6-13), eight Oil applications. All four cases are unsuccessful, since the crystals also spread to LHb. For a detailed description of the locations of the marker applications, see the text and Table 3. In the case of P1 and P9, it was impossible to differentiate between different parts of LHb, since the level of the marker application site was too caudal. Photographed in a transmitted light at a magnification of x100. Oil crystals are marked with a vertical a1Tow. Bar - 100 µm.

**Fig.3.**
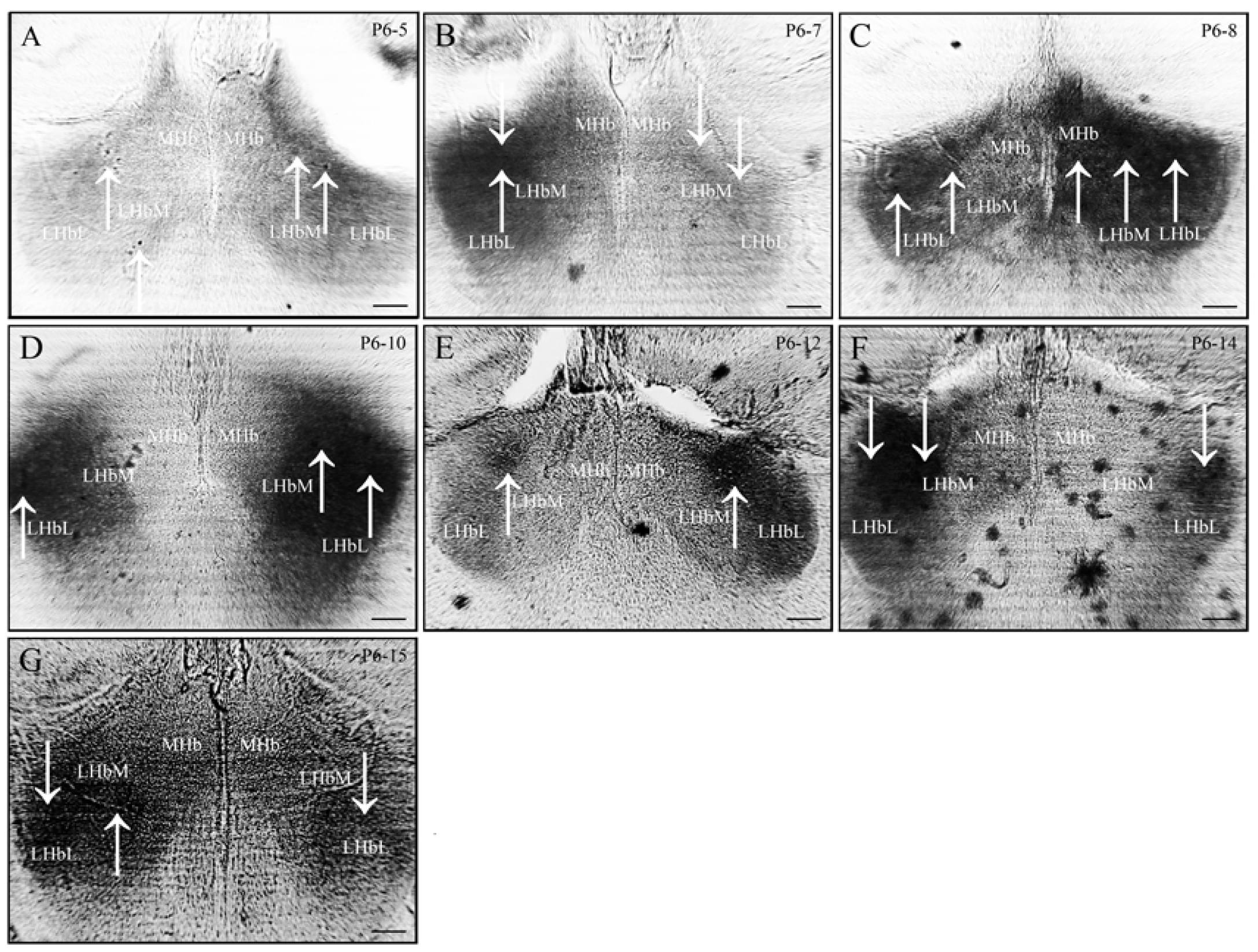
A, B, C, D, E, F, G - Bilateral places of application several Oil crystals per LHb, seven cases (P6-5, P6-7, P6-8, P6-10, P6-12, P6-14, P6-15), fourteen DiI applications. All seven cases are unsuccessful. Detailed description of places of application is available in the text and Table 4. Photographed in transmitted light at a magnification of xl00. Oil crystals are marked with a vertical arrow. Bar - 100 µm.

**Fig.4.**
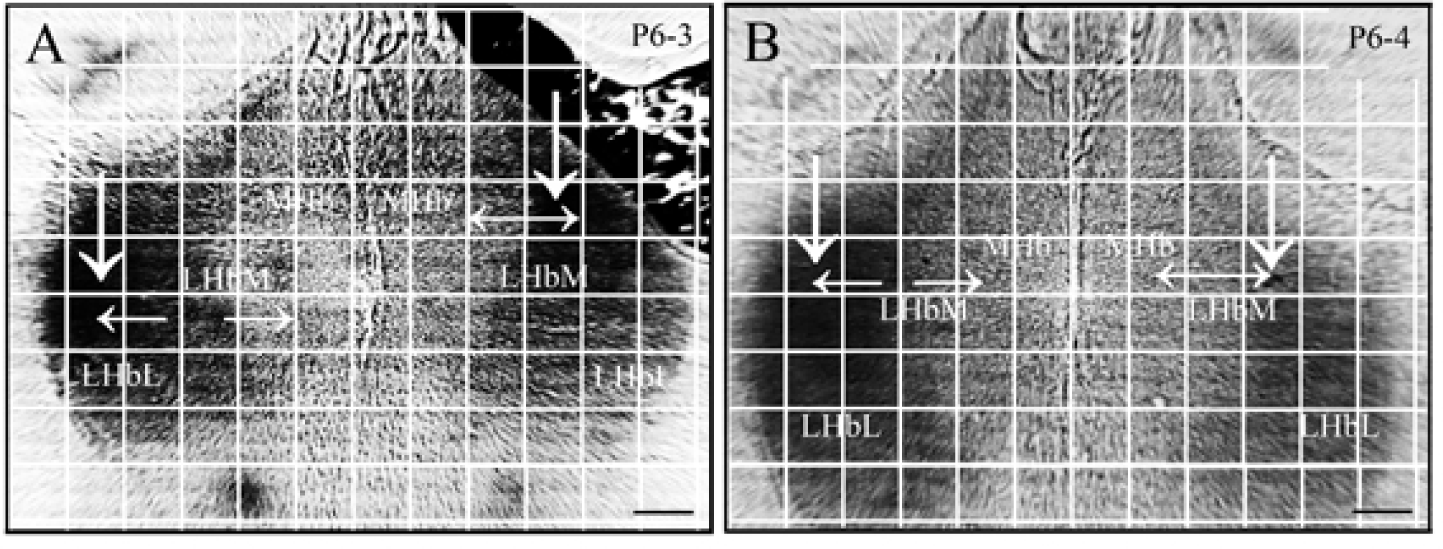
A, B - Bilateral sites for applying several Dil crystals to LHb, two cases (P6-3, P6-4), four Dil applications. Both cases are partially successful. For a detailed description of the locations of the marker applications, see the text and Table 4. Photographed in a transmitted light at a magnification of xl00. DiI crystals are marked with a vertical arrow. A coordinate grid is plotted on the image of the Hb nuclei with the site of application, showing the distance from the application site to the boundary of the Hb nuclei (Table 7). The size of one cell is 100 µm. The distance from the place of application to the boundary of the neighboring nucleus is indicated by horizontal an-ows. Bar - I00 µm.

**Table 7.** The value of the distance from the application site in the LHb DiI crystal / crystals to the neighboring MHb nucleus for successful and partially successful cases. In the completely successful case of P6-2, there are no neurons detected in TS and SFi, in some partially successful cases neurons were detected in either TS or SFi, but not simultaneously in both nuclei. The distance from the place of application to the boundary of the neighboring nucleus is taken from Pic. 4 (A; B) and Pic. 6 (A).

| Case | Type of application | Place of application (for LHb) | The distance from the application site in LHb to the adjacent MHb nucleus in one hemisphere | Success rate of the case |
| --- | --- | --- | --- | --- |
| P6-2 | Unilateral, single crystal DiI | LHbL | 350 μm | Completely successful Case |
| P6-3 on the left side | Bilateral application, several DiI crystals | LHbL | 350 μm | Partially successful case |
| P6-3 on the right side | Bilateral application, several DiI crystals | LHbM | 200 μm | Partially successful case |
| P6-4 on the left side | Bilateral application, several DiI crystals | LHbL | 300 μm | Partially successful case |
| P6-4 on the right side | Bilateral application, several DiI crystals | LHbM | 200 μm | Partially successful case |

### 3.5 Unilateral applications of a single DiI crystal - group 2 (the control group)

1. The application of a single DiI crystal on MHb is three cases (Table 5, Fig. 5A, B, Fig. 6A). All three depositions were placed exactly into the MHb nucleus. However, in two cases P6-16 and P6-17, the DiI crystal fractured during the application, and as a result, the application was not as local as planned (Fig. 5.A, B). Only in the third case of P6-11 the DiI crystal retained its integrity (Fig. 6.A). In all three cases P6-11, P6-16 and P6-17 in the adjacent MHb nucleus of the neighboring hemisphere, the marker diffusion from the application site was detected, which was especially noticeable in the smear sm marker in the neighboring hemisphere (Fig. 5D, E, Fig.6.C). In all three cases, the diffusion of the marker spread to the LHbM nucleus of the same hemisphere into which the DiI crystal was injected, even in the case of P6-11 spotting (Fig. 6.A).
2. Application of a single crystal on LHb - two cases (Table 6; Fig.5.C; Fig.6.B). In both cases, the application of a single DiI crystal was carried out exactly in the LHb nucleus. However, in case with P6-18, the DiI crystal fractured during the application to the LHbM area, which resulted in an undesirable growth of the application area. In this case, the diffusion of the marker spread to the both neighboring MHb nucleus and the MHb nucleus of the neighboring hemisphere, which is clearly visible by the sm marker of the neighboring hemisphere (Fig. 5F). In the case of P6-2, the application was carried out in the LHbL area and was the most accurate. In the neighboring MHb nucleus, there was practically no diffusion of DiI from the site of application, in the MHb nucleus of the neighboring hemisphere no diffusion was observed at all (Fig. 6D).

**Fig.5.**
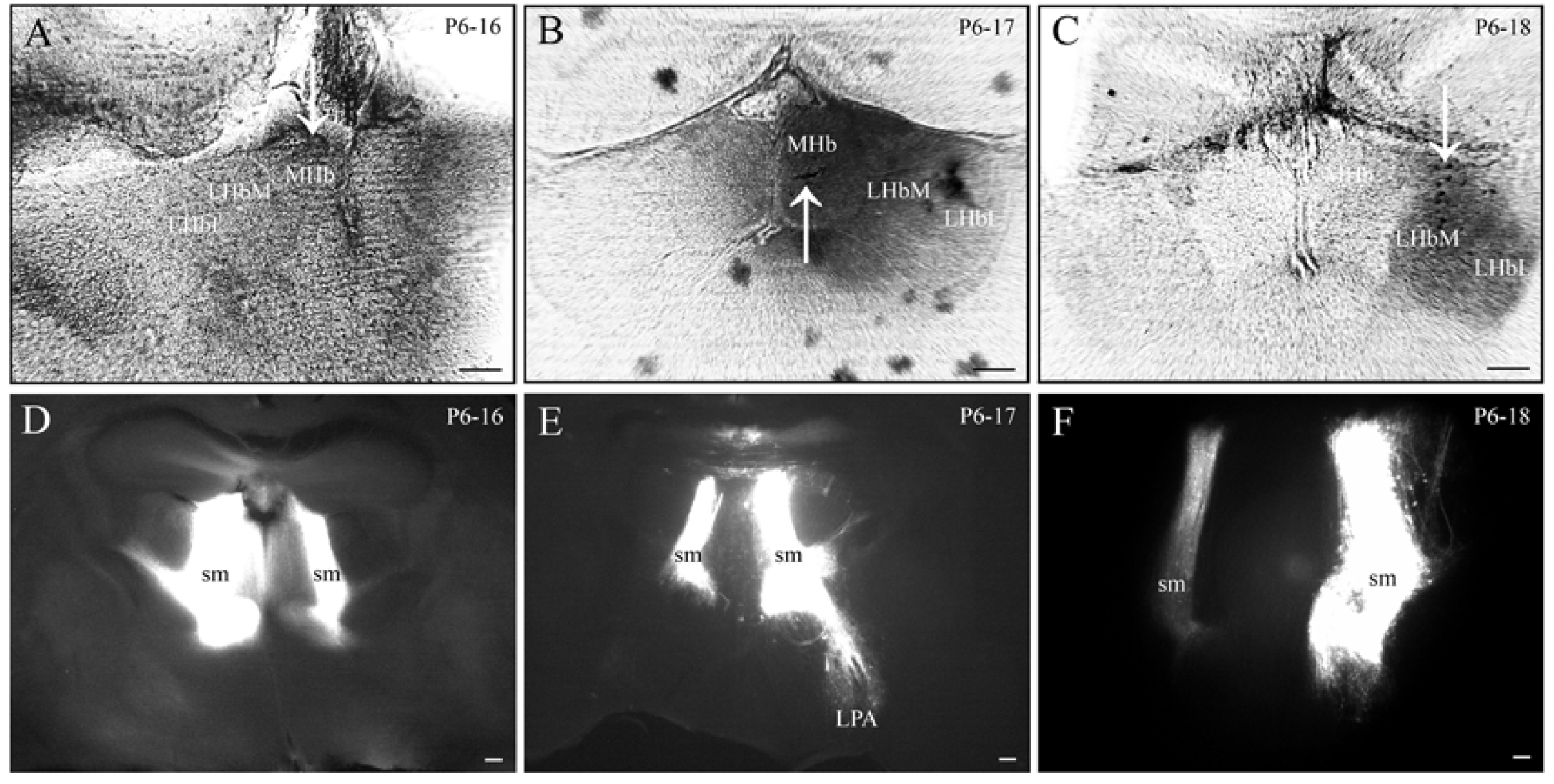
A, B - Two cases with unilateral application of a single Dil crystal to MHb (P6-16, P6-17). C - one case with unilateral application of a single Dil crystal to LHbM (P6-18). All three cases are unsuccessful; in all cases the Dil crystal was fragmented. For a detailed description of the application places of the marker, see the text and Table 5; 6. Photographed in a transmitted light at a magnification of ×l00. Dil crystals are marked with a vertical arrow. Bar - 100 µm. D, E, F is the stained Dil sm path of the neighboring hemisphere of the brain (relative to the unilateral site of the Dil crystal). D - sm path in the left hemisphere of the brain in case of P6-I 6 (Dil application in MHb on the left); E - sm path in the right hemisphere of the brain in case of P6-I7 (application of Oil to MHb on the right); F - sm path in the left hemisphere of the brain in case of P6-18 (applying Dil on LHbM on the right). Photographed in fluorescent light on an ×40 magnification. Bar - 100 µm.

**Fig.6.**
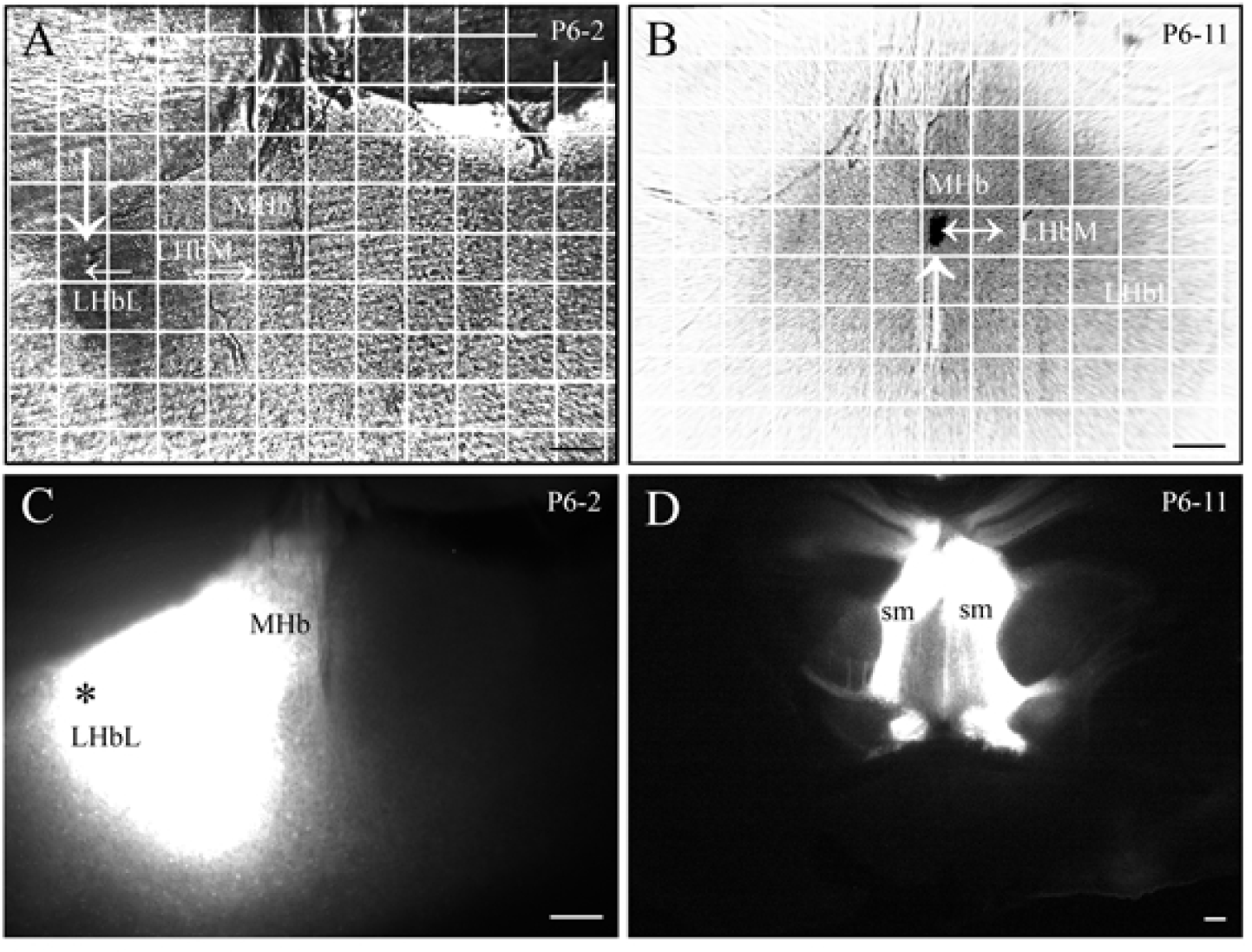
A, B - Two cases with a unilateral application site of a single Oil crystal. A - the case with application to LHbL (P6-2). B - the case with application to MHb (P6-II). Both cases are successful. For a detailed description of the places of application of the marker, see the text and Table 5; 6. Photographed in transmitted light at a magnification of ×100. A single Oil crystal is marked with a vertical arrow. A coordinate grid is plotted on the image of the Hb nuclei with the site of application, showing the distance from the application site to the boundary of the Hb nuclei (Table 7). The size of one cell is 100 µm. The distance from the place of application to the boundary of the neighboring nucleus is indicated by horizontal arrows. Bar - 100 µm. C - Unilateral site of application of a single Oil crystal on LHbL (marked with*) in the case of P6-2, photographed in a fluorescent light at an increase of ×100. Bar - 100 µm. 0 - Stained with OiI sm path of the neighboring hemisphere of the brain (with respect to the unilateral site of the Oil crystal) in the case of P6-II with the application of a single crystal in MHb. Photographed in fluorescent light on a x40 magnification. Bar - 100 µm.

### 3.6 Analysis of the presence of neurons in LPA, in TS and SFi

When in a result of marker application to LHb, neurons in TS or SFi are detected, this indicates either undesirable diffusion of the marker from the LHb nucleus into the neighboring MHb nucleus, or an inaccurate deposition. These cases are inaccurate (unsuccessful). When in a result of marker application to MHb, neurons in LPA are detected, these cases are also inaccurate (unsuccessful) for the same reasons as described above.

### 3.7 Presence of neurons in LPA (for MHb), TS and SFi (for LHb) - group 1

1. Application of several DiI crystals to MHb. All four cases with eight depositions of DiI crystals in MHb were found to be inaccurate, neuronal analysis in LPA can not be reliable, since in all cases there was a drift of crystals in LHb.
2. Application of DiI crystals on LHbM. In one application (P6-4 on the right), no neurons were detected in TS. In the case of local application of P6-3 (right), no neurons were detected in SFi. In both cases, the applications of DiI crystals were very local. These two applications are recognized as partially successful. In all other applications, neurons were detected in TS and SFi (Fig. 7A, B, Fig.8.A, B, C), and thus, they are unsuccessful.
3. Application of several DiI crystals to LHbL. In SFi, neurons were not detected in one application P6-3 (left). In one application (P6-4 on the left), in TS neurons were not detected. In both cases, the application of DiI crystals was very local. Both applications are recognized as partially successful. In all other cases, neurons were detected in TS and SFi (Fig. 7A, C, Fig.8.A), and thus, they are unsuccessful.
4. Application of several DiI crystals to LHbM + LHbL. In all six applications, neurons were detected in TS and SFi (Fig. 7C).

**Fig.7.**
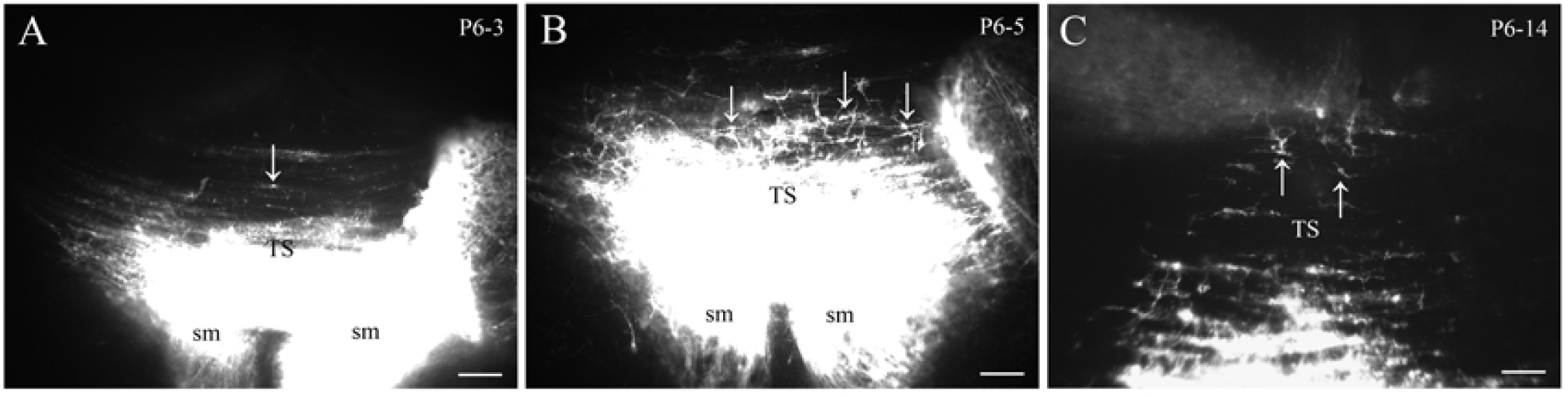
A, B, C - neurosomes stained with DiI (marked by a vertical arrow) were detected in TS after bilateral application of several DiI crystals to different parts of LHb in three arbitrarily chosen different cases: P6-3 (LHbL + LHbM), P6-5 (LHbM + LHbM), P6-l 4 (LHb + LHbL). The presence ofneurons in the TS indicates the diffusion ofthe marker beyond the boundaries of LHb nucleus. Photographed in fluorescent light. Bar - 100 µm.

**Fig.8.**
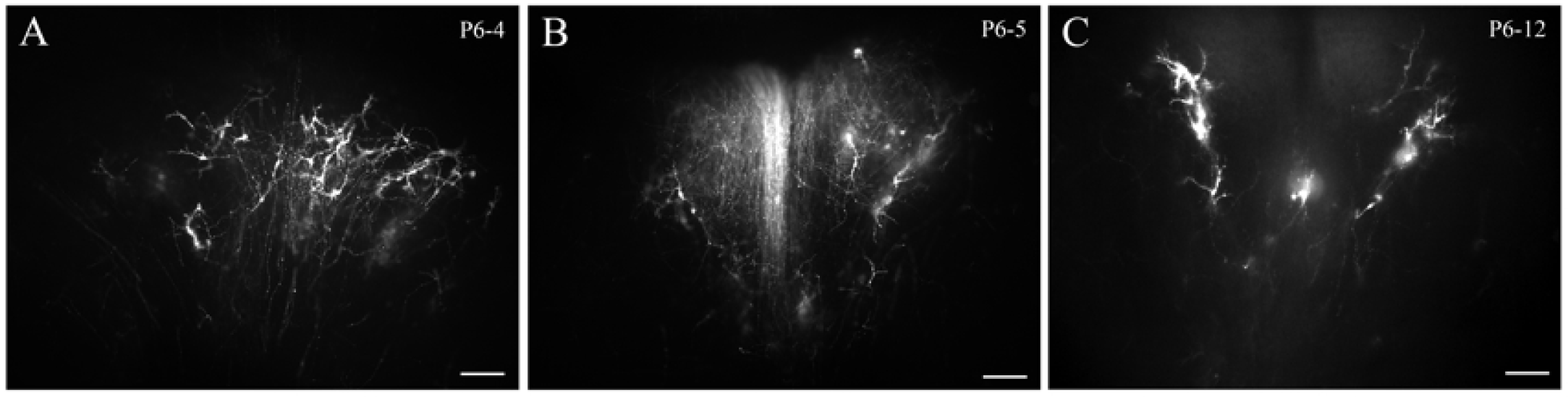
A, B, C - neurosomes stained with DiI were detected in SFi after bilateral application of several Oil crystals to different parts of LHb in three arbitrarily chosen cases: P6-4 (LHbL + LHbM), P6-5 (LHbM + LHbM), P6 -12 (LHbM + LHbM). The presence of neurons in SFi indicates the diffusion of the marker beyond the boundaries of LHb nucleus. Photographed in a fluorescent light. Bar - 100 µm.

### 3.8 The presence of neurons in LPA (for MHb), TS and SFi (for LHb) - group 2 (control group)

1. Application of a single crystal of DiI to MHb. Of the three cases with the application of a DiI crystal in MHb to LPA, neurons were detected in two cases - P6-16 and P6-17 (Pic.5E, Pic.9.A, B), in both cases, the DiI crystal fractured during the application. However, in one of these two cases - P6-16 in LPA, single neurons were detected. Thus, in accordance with the experimental scheme (Fig. 1), only the case of P11, in which no neurons were detected in the LPA, could be considered successful, in spite of the diffusion of the marker in the LHb from MHb (Fig. 6B).
2. Application of a single DiI crystal to LHbM. In the only case of P6-18 with the application of the DiI crystal to LHbM in TS and SFi, neurons were detected (Fig. 9C), including in the neighboring hemisphere of the brain, but the DiI crystal was fractured during application, which undesirably increased the area of application, and the diffusion of DiI spread to the neighboring nucleus of the MHb of the neighboring hemisphere. This is an unsuccessful case.
3. Application of a single crystal of DiI to LHbL. In the single case P6-2 with DiI crystal application to LHbL in TS and SFi, neurons were not detected. The DiI crystal retained its integrity during application. This is a completely successful case.

**Fig.9.**
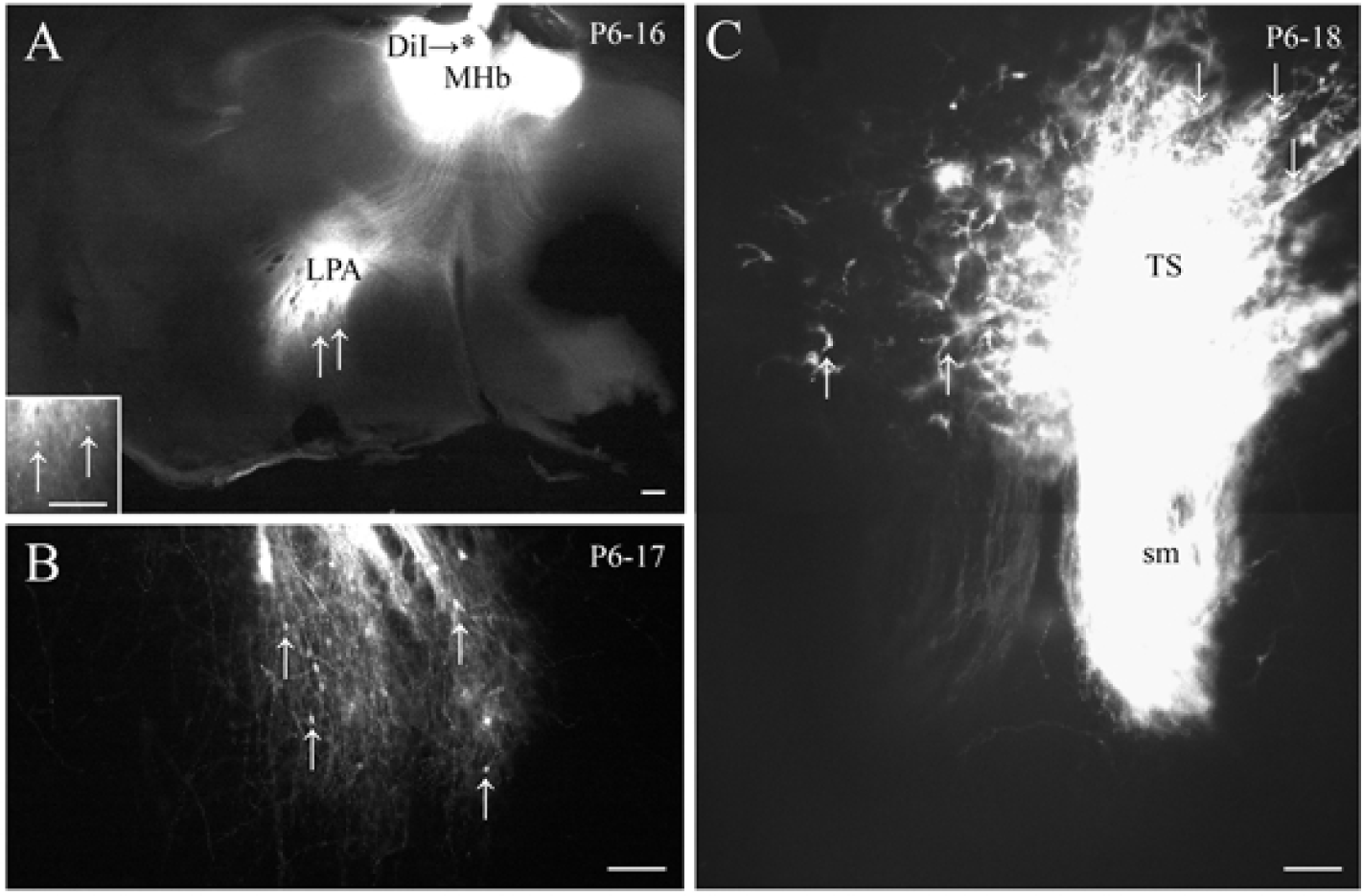
A, B - stained with Oil neurosomes (marked by a vertical arrow), revealed in LPA after unilateral application of a single Oil crystal in MHb (the place of application is marked with *), the Oil crystal was fragmented during application. The presence of neurons in the LPA indicates the diffusion of the marker beyond the boundaries of MHb nucleus. Photographed in a fluorescent light at x40 and xl00 (inset). Bar - l00 µm. C - neurosomes stained with DiI were detected in TS after unilateral application of a single Oil crystal in LHbM, the DiI crystal fractured during the application. The presence of neurons in the TS indicates the diffusion of the marker beyond the boundaries ofLHbM nucleus. Photographed in a fluorescent light at xl00 (inset). Bar - 100 µm.

## 4 DISCUSSION

### 4.1 The problem of using the DiI method for higher vertebrates nerve connections research

Currently, studies of the neural connections of higher vertebrates with the DiI method are practically not being carried out, this method was the most popular in the 1990s. This is mainly related to shortcomings of this method, in particular the inability of DiI to dye the fully formed myelinated neural tracts of adult specimens (Molnar Z, Blakemore C, 1995), while the vast majority of neuroanatomical work was done particularly on adult animals. For this reason, the bulk of work with DiI on higher vertebrates was performed on embryos and fetuses, in which the neural tracts have not been myelinated yet. The main purpose of this series of studies was to study the formation of neural connections of specific brain structures at different stages of embryonic development (Catalano S, Robertson R, Killackey H. 1991; Linke R; Frotscher M. 1993; Kitzes L et al. 1995).This is a very narrow and specialized topic, represented by single isolated works. An additional complication is the fact that it is impossible to clearly differentiate the axons and dendrites of neurons dyed with DiI, and this considerably complicates the interpretation of the data obtained. However, DiI and its analogues are currently the only markers that generally allow one to investigate the development of neuronal connections of higher vertebrates, and the possibility to completely eliminate the use of stereotaxis in DiI studies does not allow us to completely abandon the use of this method, despite all its shortcomings(Table1).

In a result of my previous preconceptual studies of Hb connections in rats at different stages of embryonic and postnatal development (Klepukov A, Makarenko IG, 2013) two significant problems were discovered rather quickly, which are usually not emphasized. The first problem is the inaccuracy of DiI application and the extensive diffusion of the marker at the place of application of the marker, which makes it impossible to clearly distinguish the connections of the individual nuclei of the studied brain structures to which DiI applications were made, especially if DiI applications were made on the sagittal section of the brain. The second problem, which is the consequence of the first one, is a false positive result.

Solving these two problems, at least partially, as well as increasing the percentage of successful applications with reliable results (efficiency of the method) is the main goal of this new work.

### 4.2 The problem of the accuracy of applying DiI to the desired structure of the brain

Despite the fact that most of the work with DiI has been devoted to the study of the formation of neural connections of various brain structures in embryos and fetuses of rats or mice, none of these studies raised as the main question how accurate the detection of DiI of the corresponding nerve connections is. Nevertheless, the fact that at present time the work with DiI is practically scaled down, suggests that this problem in combination with others is the key problem. In all the works I have studied (Catalano S, Robertson R, Killackey H. 1991, Linke R, Frotscher M. 1993, Kitzes L et al., 1995) it was indicated in the methodical part that DiI crystals / crystals were applied manually to the brain slice with the aid of a thin glass capillary. This approach to application has three significant drawbacks. The first drawback is that none of the studied research focused on how exactly the number of DiI crystals affects the accuracy of application, mainly variations were related to the aggregate state in which the marker was introduced, for example in the form of a crystal or liquid (von Bartheld CC, Cunningham DE, Rubel EV 1990). In some studies it was pointed out that a single DiI crystal was applied (Kil J et al., 1995), but no conclusions were drawn from this. The second drawback is the technique of applying DiI crystals / crystals. In none of the works that I analyzed was not specified whether the DiM micromanipulator was used for DiI application, in all the works it was clearly indicated that the DiI crystals were applied manually, and this a priori can not guarantee a stable and accurate depth of penetration into the tissue. In my own preliminary study (A. Klepukov, I. Makarenko, 2013) a significant number of cases were eliminated because of poor-quality manual application of DiI crystals, as a result two Hb nuclei were widely labeled instead of one. The third drawback is the application of DiI crystals / crystals to a conditionally arbitrary slice of the brain.

Based on the analysis of the literature, it is possible to make a conclusion that the brain slice is almost always made on the basis of very approximate assumptions about the correspondence between the level of the desired structure and the level of the slice. None of the studies indicated how successful the section of the brain with the desired structure was, and how many cases were eliminated as a result of unsuccessful sections. In my own preliminary study, a significant number of cases were lost even before the application of DiI crystals, since the section was always made blindly.

A partial solution of these three methodological problems with the application of DiI is presented in this paper.

The first problem - the estimation of the effect of the amount of a marker on the extent of diffusion at the application site - is partially solved by reducing the number of DiI crystals from several to one. The results of applying several DiI crystals to MHb are very indicative, since no completely successful case with precise application has been obtained in which diffusion of the marker would be limited to only one nucleus. In the case of the applycation of several crystals on LHbM or LHbL, the number of successful cases estimated by the absence of neurons in structures that did not innervate the nucleus with the marker was also zero, although there were partially successful cases (P6-3, P6-4). The more extensive application was, the lower is the percentage of partially successful cases. Nevertheless, even when a single DiI crystal was applied to a single Hb nucleus, it was not possible to completely avoid unwanted diffusion into neighboring nuclei. Moreover, in all three cases, when a single DiI crystal was applied to MHb (P6-11, P6-16, P6-17) in the MHb nucleus of the neighboring hemisphere, an extensive diffusion of the marker was also recorded, up to the dyeing of the neighboring path - sm (fig.5.DE; fig.6.D). A new unexpected problem, which was revealed when a single crystal was applied to a tissue, was that the crystal could be fractioned directly at the time of application. This resulted in an undesirable growth in the application area and, as a consequence, a wider marker diffusion and loss of accuracy (cases P6-16, P6-17, P6-18). As the result, only two cases out of five (unilateral applications) can be considered conditionally successful in the accuracy of application aspect - P6-2 and P6-11. In both of these cases, a single DiI crystal was applied exactly to one of the Hb nuclei, and the crystal did not break up into parts (Fig. 6.A, B). In the case of P6-2, no neurons were detected in TS and SFi when applying the marker in LHbL, neurons were not detected in the case of P6-11 when the marker was placed in MHb in LPA. In the sense of the absence of detected neurons in structures that do not innervate the nucleus with the marker, these two cases can be considered completely successful.

The second problem - manual application of the marker with the help of a capillary - can be easily solved with a micromanipulator (Narishige C-1). However, despite the fact that the accuracy of the introduction of a glass needle with crystals / crystal increases significantly, there is a number of completely unsolvable problems that have clearly been revealed in this study. It was very difficult to control the size of the application when applying several DiI crystals, for this reason some cases were lost (P1-application left, P9-application left), because the application was too extensive and covered both nuclei simultaneously. Sometimes there was another problem, and though the main DiI crystals were applied accurately, but the minor part (dust from DiI crystals) spread on the neighboring nuclei, which was also undesirable and led to loss of the case: P1-right application (Pic. 2A), P6 is the application to the right (Pic. 2B), P9 is the application to the right (Pic. 2C). When a single crystal was applied, the problem of crystals spreading to adjacent structures did not exist, but a single DiI crystal was often fractured at the time of application, which resulted in an undesirable extension of the application area. Even if a single DiI crystal did not break up into pieces, the size of the application was still difficult to control, since it still directly depended on the size of originally chosen DiI crystal, and the choice of a DiI crystal with a given size was very difficult. Finally, despite the use of a micromanipulator to control the depth of application, this did not help in the case of application of single crystals, which were not always firmly fixed in the tissue, leading to a loss of cases (P6-19, P6-20). In this case, even covering the section area with a layer of agar (analogue of the paste fixing the crystal at the place of application) was not always effective, because the DiI crystal could still fall away from the place of application to the neighboring brain structures at the time of agar application.

The third problem - the application of DiI crystals/crystal to the precise slice of the whole brain - can be solved with the help of controlled sequential slicing of the brain on a vibroslicer with parallel analysis of each section for the identification of the necessary structures. This method takes more time than a single random slice of the brain at the conventional level of the desired structure, but the method of consecutive slicing is much more accurate. However, even at this stage, some losses are possible, in particular, in two cases (P6-1, P6-9), the level of the section was too caudal (Fig. 2A, Fig. 2C), which was found only when the brain with the already marked marker was completely cut into sections and analyzed. A possible solution of the problem of capturing the desired section, which detects the correct level of the brain with the desired structure, would be a consistent contrast staining of the sections (for example, with methylene blue). On a contrasting colored section, it is easier to reveal the desired structure and consequently the brain level.

### 4.3 The problem of choosing the stage of development of the object of research and the false positive result

The result of extensive diffusion or inaccurate deposition of DiI crystals / crystal is a false positive result, meaning that in structures of the brain that do not innervate the nucleus with the marker (according to literature data), stained neurosomes or fibers that should not be present in these brain structures are detected.

The problem of a false positive result is solvable if the results that are compared are received at least with use of two different independent methods. The coincidence of the results obtained by different methods indicates their reliability. The agreement of results obtained by different methods indicates their reliability. However, in the case of DiI during the study of the formation of neural connections at different times of embryonic and neonatal development it is hardly possible, since DiI and its analogs are the only markers that can detect neural connections specifically at early embryonic and neonatal stages. Stereotaxis as an alternative method with other markers different from DiI, especially on embryonic stages, is not appropriate.

Nevertheless, the problem of comparing the results obtained by different methods can be circumvented by using in the study animals on the stage of development when all the main neural tracts have already been formed. A comparison of already formed neural connections will make it possible to understand how accurate the results obtained with both DiI and stereotaxis are. Although in this case we compare the data obtained at different stages of development (since DiI does not work for adult neural tissue), but if the main tracts and connections are already formed just before the brain enters the active phase of myelination, then such a comparison is permissible.

My own preliminary research (A. Klepukov, I. Makarenko, 2013) allows suggesting that in rats at least on the 6th day of postnatal development the main conducting systems of Hb are already formed. Literature data reveals that by the 7th day of postnatal development in rats and mice the myelination processes in different parts of the central nervous system already begins (Jacobson S, 1963, Banik N, Smith M, 1977; Rozeik C, Von-Keyserlingk D, 1987).

As far as is known, DiI does not work on myelinated tissue (Molnar Z, Blakemore C, 1995). Thus, the upper age limit for neonatal rats and mice, on which DiI can still be used, is approximately the 6th day of postnatal development. By this time, all the main links of Hb with other structures should already be formed. Originally this work was planned as a test of this particular assumption.

The scheme of the connections of different Hb nuclei with forebrain structures is described in details in the literature both for rats (Herkenham M, Nauta WJ, 1977) and partially for mice (Qin C, Luo M, 2009; Broms J et al., 2017). It is known that LHb is connected by afferent one-sided connections with LPA, MHb is connected by afferent one-way connections with TS and SFi. The higher the percentage of cases with the application of a marker to the corresponding nucleus fit into such a scheme (Fig. 1), the higher is the overall efficiency of the method and the lower is the percentage of cases with a false positive result.

Among the cases with bilateral application of several DiI crystals, the neuronal distribution in none of the twenty-six (0%) applications in TS and SFi nuclei was consistent with what was described in the literature. For cases with unilateral application, two applications out of seven were already coincided with the literature data (28.57%, or 40%, except for two cases of P6-19, P6-20 that were not analyzed, since the DiI crystal did not become fixed in the tissue) while in both successful applications the DiI crystal retained its integrity. Despite the fact that samples of different sizes are compared, the trend is obvious. However, the only one case can be considered quite successful: P6-2 with DiI deposition in LHbL, since in the case of P6-11 when DiI was applied to MHb, the diffusion of the marker spread to the MHb of the neighboring hemisphere, and to the neighboring LHb nucleus, although in LPA neurons did not come to light (there is no explanation for this yet), so this is a formally successful case if we follow the logic of the experiment (Fig. 1).

A single non-fragmented DiI crystal gives a lower diffusion level at the application site and as a consequence a lower percentage of cases with a false positive result. Ideally, if a method of applying DiI to Hb (or to any other well-studied brain structure) is developed, 95% of the depositions yield a reliable result (coinciding with the description in the literature), then this technique can be considered successful and suitable for reliable research of brain connections that are not described yet. The development of such a technique is the topic of the next work.

### 4.4 The problem of determining the DiI diffusion boundaries

Unilateral applications of DiI showed significantly higher efficacy (28.57% vs. 0%) with a lower sample range (7 applications versus 26). However, only the application of DiI (P6-2), which was done at the level of the lateral border of LHbL, was completely successful, while the distance from the application site to the lateral border of MHb was 350 µm (Fig. 6A, Table 7).

The case of P6-11 is only formally successful, since in the LPA, despite the apparent diffusion of a marker from MHb to LHb, neurons have not been found and there is no explanation for this yet, so for this particular case, the importance of distance value from the place of deposition to the neighboring nucleus (150 µm) is conditional. The MHb diameter at the P6 term does not exceed 200 µm (Paxinos et al., 2007). In all three cases, with the unilateral application of the DiI crystal in the neighboring hemisphere, the sm path was stained (Fig. 5D, E; Fig.6.D), which indicates the expansion of marker diffusion beyond the MHb nucleus irrespective of whether the crystal is fractured or not. In other words, if the nucleus size is less than 200 µm, the probability of inaccurate application is high in any case. However, this figure (200 µm) needs further clarification.

Despite the fact that cases with bilateral deposition of several DiI crystals on the LHb were not generally successful, there were also partially successful cases (P6-3, P6-4). Partially successful is the case in which neurons were detected either in TS or SFi, but not in both nuclei simultaneously (Table 4). In both partially successful cases, the distance from the place of deposition to the boundary of the neighboring nucleus was not less than 200 µm (Fig. 4A, B, Table 7). For successful applications, the size of the DiI application and the size of the brain nucleus in which the DiI application is performed are important. Probably the size of the brain nucleus should be no less than the distance from the lateral boundary of LHbL to the lateral boundary of MHb multiplied by 2 if the marker migrates equally easy in all directions from the site of application, which corresponds to 350×2 = 700 µm (calculated for the completely successful case of P6-2). It is also necessary to take into account additional factors, for example, the presence of a capsule of nerve fibers around the nucleus, which can prevent the marker from diffusing out of the nucleus (that is why for LHb, the range of diffusion of the marker is still limited to 350 µm).

In the case where the distance between injections in the bilateral application of DiI on the Hb nuclei is less than 350 µm, DiI diffusion fluxes from the application sites may occur if the distance between the DiI application sites is small (for more laterally located brain structures, the percentage of successful cases may be higher).

However, the refinement of the diffusion boundaries of the marker requires further investigation, and the resulting figure for the permissible diffusion boundaries of DiI at the application site (700 µm) is not ultimate, so it cannot be used as a final conclusion.

### 4.5 The problem of alternative control cases

A particular problem is the fact that even for Hb, which connections are well described for rats, there is still no complete scheme of afferent connections for mice described in the literature, despite a number of qualitative studies already done (Qin C, Luo M, 2009; Broms J et al., 2017). In other words, there is some (very low) probability that mice’s MHb can be associated with LPA, and mice’s LHb with TS and SFi. Corresponding applications of DiI to LPA could disprove such a hypothetical scheme of innervation, TS and SFi (control cases), thanks to which it would be possible to check the distribution of labeled nerve endings already in Hb itself, but given the excessive degree of DiI diffusion, this task is currently unfeasible for a number of reasons.

It is known that, in addition to TS and SFi with MHb, it is also associated with afferent connections of Bed nucleus of stria terminalis (BST), which is adjacent to the ventral surface of TS and SFi. BST is also associated with afferent connections to LHb (Herkenham M, Nauta WJ, 1977). When DiI is applied to the TS in the event of the slightest hit of the marker due to diffusion into the BST, there is a high probability of obtaining a false positive result in LHb. The same applies to LPA, the dorsal boundary that also bounds to BST, as a consequence, in the case of diffusion of a marker from LPA to BST, it is possible to get a false positive result in MHb. Thus, control cases on other structures of the brain can hardly help clarify the picture of the true innervation of different Hb nuclei.

## 5 CONCLUSION

1. To obtain the most reliable results in the study of brain connections, it is preferable to use unilateral applications of single crystals, since their accuracy is generally higher (28.57%) than for bilateral applications of several crystals (0%).
2. To obtain accurate application of DiI to the desired brain nucleus, it is necessary to use a micromanipulator, which allows controlling the depth of application. The application should be carried out strictly on the section of the whole brain, exactly corresponding to the level of the brain structure being studied. The exact section of the whole brain should be detected with the help of a sequential controlled under a stereomicroscope slicing the brain on a vibroslicer with a parallel analysis of each section to identify the necessary structures. Slicing the whole brain should stop exactly at the exact moment when the desired brain structure is accurately identified.
3. When applied to the desired brain structure, a single DiI crystal is often fractured during the application, which leads to an undesirable extension of the application area. Even if the single DiI crystal is not fractured, it is still difficult to control the degree of undesired diffusion, since it still directly depends on the size of initially chosen DiI crystal, and the size of the DiI crystal is a poorly controlled value. Given this, the overall accuracy of the DiI injection method, even with use of a single crystal, remains low.

## 6 ACKNOWLEDGMENTS

The author of the work (Klepukov AA) expresses gratitude to Lovatu M.L (the head of the Scientific Research Institute of Mitineniring of the Moscow State University) for conducting the stage of perfusion and euthanasia of neonatal mice. The author of work (Klepukov A) also expresses his gratitud to Ekaterina Georgievskaya and Ekaterina Antipova for translating the whole text into English.

This private initiative fundamental research work was carried out exclusively on 100% personal funds of the leading embryologist of the clinic Altra-Vita Klepukova A. in accordance with the internal program of the clinic to improve the professional level of employees.

## 7 AUTHOR CONTRIBUTION

Klepukov A: author of the concept and article, the performer of the experimental part of the work 98%. Sponsor of research on 100%. Apryshko V.P: participation in the work is to ensure delivery of biomaterial from the Biological Faculty of Moscow State University (the service is provided only to MSU employees). Yakovenko S.A: participation in the work is the registration of work in the state register of the EGISU www.rosrid.ru, registration number AAAA-A17-117111440045-7 (the service is provided only in the name of the director of the organization, necessary in Russia).

